# Vitamin C prevents *S. aureus* infections of endothelial cells through its interaction with specific residues in the active site of gC1qR/p33 involved in direct entry, notably SER230, ASN345, ILE348, TYR224, ASN346, SER347, and GLU438, and its potential inhibitory effects on FnBPs, α5β1, and Protein A molecules

**DOI:** 10.1101/2024.03.27.586894

**Authors:** Maroua Miliani, Zeyneb Hadjidj, Zakariya Bensefia, Sara Dahou, Nasreddine Ammouche, Imane Hamdoun, Hadjer Benammar, Sarah Benammar, Mourad Aribi

## Abstract

*Staphylococcus aureus* (*S. aureus*), often perceived as an extracellular pathogenic agent, exhibits a remarkable ability to penetrate host cells, including endothelial, epithelial, and osteoblastic cells, significantly contributing to the pathogenesis of infections. A significant pathway for this invasion appears to involve the bacterium’s binding to the α5β1 integrin *via* a fibronectin bridge, followed by phagocytosis. Additionally, *S. aureus* presents staphylococcal protein A, a cell wall protein that binds to the Fc and Fab regions of immunoglobulins, playing a crucial role in virulence and immune evasion, and can also bind directly to the gC1qR receptor on endothelial cells. Furthermore, vitamin C is recognized for its antimicrobial and immunomodulatory properties, offering potential for reducing the risk of infection. Considering the aforementioned elements, our study focused on exploring the potential effects of vitamin C on the interactions between *S. aureus* and endothelial cells. Thus, we particularly examined two aspects: on the one hand, interactions involving fibronectin-binding proteins (FnBPs) proteins and human α5β1 integrin, and on the other hand, the interaction between vitamin C and the direct entry receptor (the globular heads of complement component 1q receptor [gC1qR/p33], also known as hyaluronic acid binding protein 1 [HABP1]). To achieve this, we utilized molecular modeling assays, primarily relying on molecular docking.

## 1. Introduction

*S. aureus* is an opportunistic pathogen capable of colonizing various cellular niches, endowing it with high adaptability and virulence, which can lead to various diseases in humans when it enters the bloodstream. Its ability to live inside host cells plays a crucial role in infection progression. Endovascular infections caused by *S. aureus*, such as infective endocarditis (IE), are associated with high morbidity and mortality rates. The molecular mechanisms underlying *S. aureus* colonization and tissue invasion are complex. Experimental data show that plasma proteins, endothelial cells, and components of the subendothelial matrix play an important role in the pathogenesis of endovascular infections caused by *S. aureus*. Bacterial adhesion to host cells and tissues is a crucial step in its pathogenesis, and it has become evident that many adhesins recognize more than one ligand. For example, it has been demonstrated that Staphylococcal protein A (SpA) binds to von Willebrand factor and another ubiquitously expressed cellular protein, designated gC1qR/p33 (gC1qR). gC1qR is highly expressed on activated platelets and endothelial cells [1]. Its virulence is indeed attributable to the presence of surface proteins, including protein A, which promotes immune system evasion by binding to the gC1qR receptor on endothelial cells.

Importantly, vitamin C (L-ascorbic acid), is one of the most common and essential vitamins. Additionally, beyond its protective role and ongoing exploration of novel functions, it stands as one of the most cost-effective treatment options capable of enhancing immune function, alleviating allergic reactions, fighting infections, and potentially inhibiting cancer development [2,3]. Although vitamin C itself is a potent antioxidant, its aerobic metabolism increases oxidative stress on bacterial cells [4]. Thus, vitamin C can be a safe and natural alternative to limit the growth of *S. aureus* when non-toxicity is required [5]. Moreover, in addition to its well-known immunomodulatory effects, it has recently been shown to possess immunomodulatory properties opposing interesting endothelial cell infection [6].

Molecular modeling has become a new technique for understanding chemical phenomena and a working tool in various fields other than structural chemistry, such as medicine, cell biology, biochemistry, immunology, etc., as well as in the calculation of potential energy surfaces of organic molecules [7,8]. Among the molecular modeling methods are docking and molecular dynamics (MD) [9,10]. These computational tools are very useful in biology, pharmacy, and medicine, as most active ingredients are small molecules (ligands) that interact with a biologically relevant target, usually a protein (receptor).

Considering aforementioned above, our objective was to employ molecular modeling methodologies to assess the potential impact of vitamin C on the interactions between *S. aureus* and human endothelial cells. This included examining the involvement of Staphylococcal protein A, FnBP proteins and human α5β1 integrin, as well as studying the interactions between vitamin C and the active site of the direct entry receptor gC1qR/p33.

## 2. Materials and Methods

Our aims encompassed predicting the interactions between vitamin C and the target proteins FnBPs and α5β1 staphylococcal, examining the existing interactions between vitamin C and the binding site of the direct entry receptor gC1qR/p33 of *S. aureus*, and investigating its impact on the interaction between Staphylococcal Protein A and endothelial cells. Protein structures and ligands were obtained from the Protein Data Bank (PDB) [11] and PubChem databases [12]. Molecular docking was performed using ArgusLab software [13], with subsequent visualization of interactions carried out via Discovery Studio [14]. The steps of our virtual screening included protein and ligand selection and preparation, molecular docking, and post-docking analysis.:

### 2.1. Microcomputers

We utilized a microcomputer with 16GB memory, an AMD Ryzen 5 3400g CPU 3.5GHZ processor, and Windows 11 operating system for studying the effects of vitamin C on interactions with FnBPs and α5β1. For exploring interactions between vitamin C and the direct entry receptor gc1qr/p33 of *S. aureus*, as well as studying the effect of vitamin C on staphylococcal protein A, we used a microcomputer equipped with 3GB RAM, an Intel(R) Core i5-480M processor clocked at 2.66 GHz with 3 MB cache, and Windows 7 version 2007 operating system.

### 2.2. Tools used for computational study

Several programs were employed for this work, including ArgusLab for molecular docking, which is based on quantum mechanics for electronic structure prediction, ChemSketch for designing 2D or 3D complex structures and modifying chemical structure images, Chem3D for visualizing and sharing three-dimensional (3D) structures of molecules, and Discovery Studio, a software suite used for simulating small and macromolecular systems, including simulation, ligand design, and pharmacophore modeling. It was used to visualize poses and interactions between vitamin C and protein active sites.

### 2.3. Databases

We utilized the PDB as the global archive for structural data of biological macromolecules, providing 3D structures of biological macromolecules, including proteins and nucleic acids obtained through various methods such as X-ray crystallography and NMR spectroscopy. Additionally, PubChem served as the American database of chemical molecules managed by NCBI, containing small molecules as well as larger molecules such as nucleotides, carbohydrates, and peptides.

### 2.4. Protein preparation

The three-dimensional structure of the FnBPs protein was downloaded from the Protein Data Bank (PDB), with the PDBID “1FNF” corresponding to the human fibronectin fragment including repetitions 7 to 10 of type III. The crystallographic structure of the gC1qR/P33 protein was obtained from the PDB under the code PDB: 6szw, and the 3D structure of the protein was examined and prepared using ArgusLab. The active site was identified, and water molecules and other ligands were removed. The three-dimensional structure of the SpA protein was downloaded from the PDB under the code PDBID “4WWI". This crystalline structure of the C domain of the staphylococcal protein A in complex with the FC fragment of human IGG was resolved by X-ray crystallography with a resolution of 2.3 Angstroms.

### 2.5. Ligand preparation

The ligand, ascorbic acid, was downloaded from the PubChem platform in 2D form, and its chemical structure was modeled using ChemSketch software. The ligand’s availability was verified via the Molinspiration server.

### 2.6. Molecular docking

Molecular docking was performed to assess the effect of vitamin C on (i) FnBPs and α5β1 Staphylococcal molecules using a three-dimensional grid encompassing the active site, (ii) intermolecular interactions between vitamin C and the active site of the direct entry receptor gc1qr/p33 of *S. aureus*, and (iii) interactions between vitamin C and staphylococcal protein A, using crystalline structures of the C domain of this protein. The protein A was embedded in a three-dimensional grid largely encompassing the active site, with a center defined by coordinates (X=20Å, Y=23Å, Z=21Å). Ligand-protein interactions were visualized using Discovery Studio.

## 3. Results and Discussion

### 3.1. Impact of vitamin C on S. aureus-FnBPs, α5β1-endothelial cell interactions

#### 3.1.1. Insights from molecular docking analysis

The configuration of binding sites, spatial delineation of interaction zones, and dynamics of molecular interactions between vitamin C and the FnBPs protein are respectively depicted in Figures 1, 2, and 3.

**Fig. 1.**
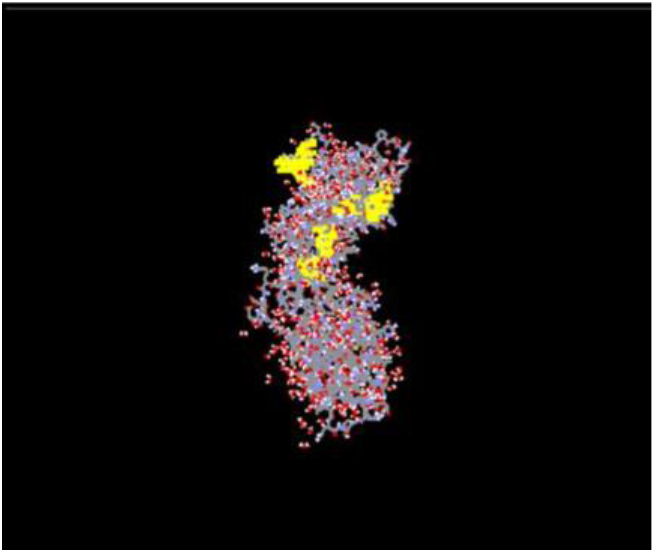
Binding site on ArgusLab. The program provided us with 150 poses, and we selected pose 1, which was the most significant with an energy of -6.56802 kcal/mol.

**Figure 2.**
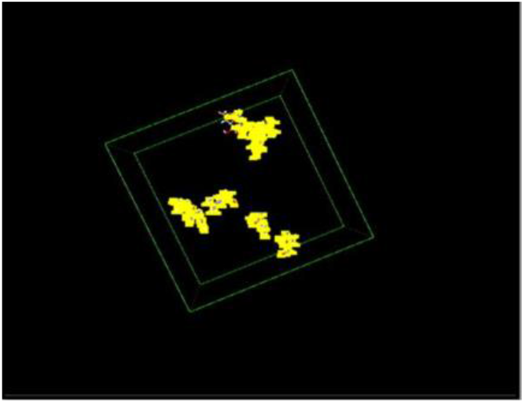
Binding sites.

**Fig. 3.**
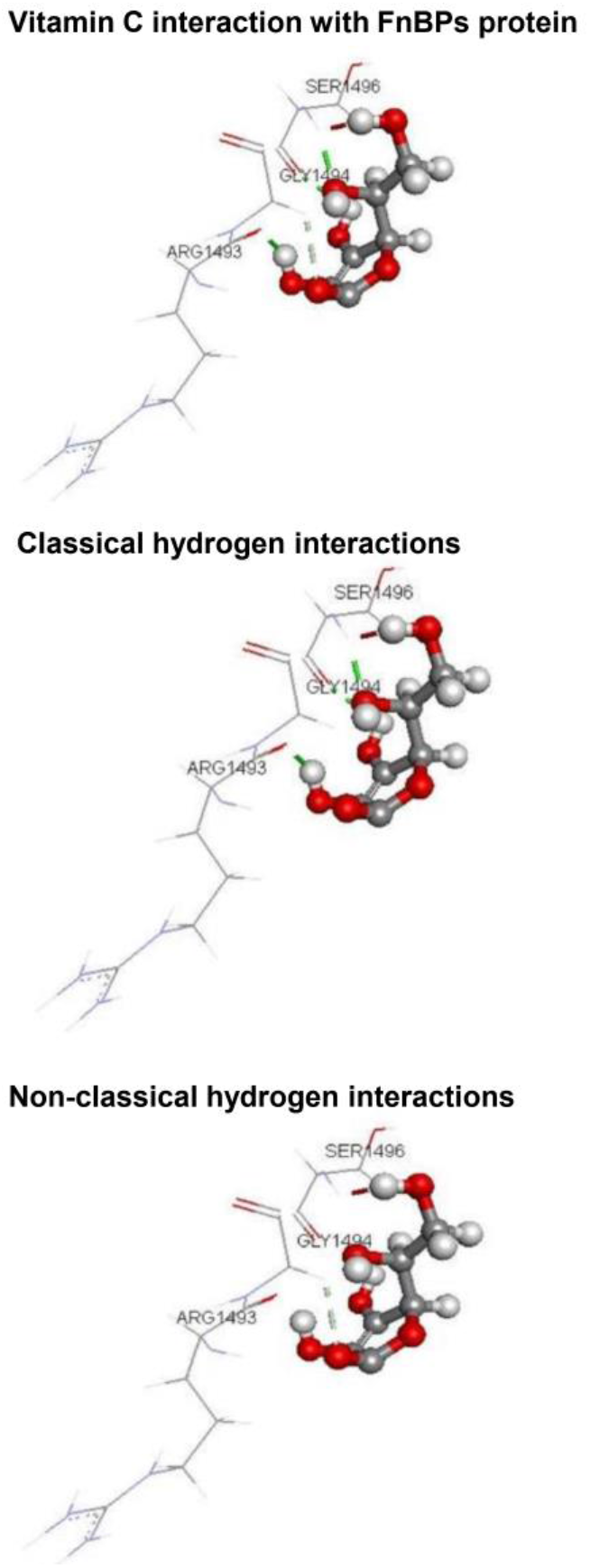
Interaction between vitamin C and the FnBPs protein.

The obtained results indicate that vitamin C meets Lipinski’s criteria, suggesting adequate bioavailability and potential favorable interaction with target proteins. Molecular docking analysis reveals several plausible poses, with the most favorable showing significant interactions with residues of the FnBPs protein’s active site. Moreover, the interaction between vitamin C and the FnBPs protein reveals the formation of hydrogen bonds, suggesting a possible inhibitory effect on bacterial entry into endothelial cells. Complex 1FNF with the lowest Score energy (−6.56802 kcal/mol) formed 4 interactions with our ligand. Hydrogen interactions are formed with the following amino acids: SER1496:HN with a distance of 1.98Å, SER1496:O with a distance of 2.19 Å, GLY1494:HA1 with a distance of 2.37 Å, as well as ARG1493:O with a distance of 1.84 Å (Table 1), suggesting that vitamin C could have an effect on the staphylococcal protein A. The top 10 results, detailing the ligand poses, are presented in Table 2. We determined the dimensions of the active site bounding box to be X = 40, Y = 40, Z = 60, as shown in Table 3.

**Table 1.** Types of bonds formed between the protein and the ligand.

| Molecule | Hydrogen interaction |  |
| --- | --- | --- |
|  | Involved residues | Distance (Å) |
| Ascorbic acid | SER1496:HN | 1.98112 |
|  | SER1496:O | 2.19359 |
|  | GLY1494:HA1 | 2.37902 |
|  | ARG1493:O | 1.84572 |

**Table 2.**
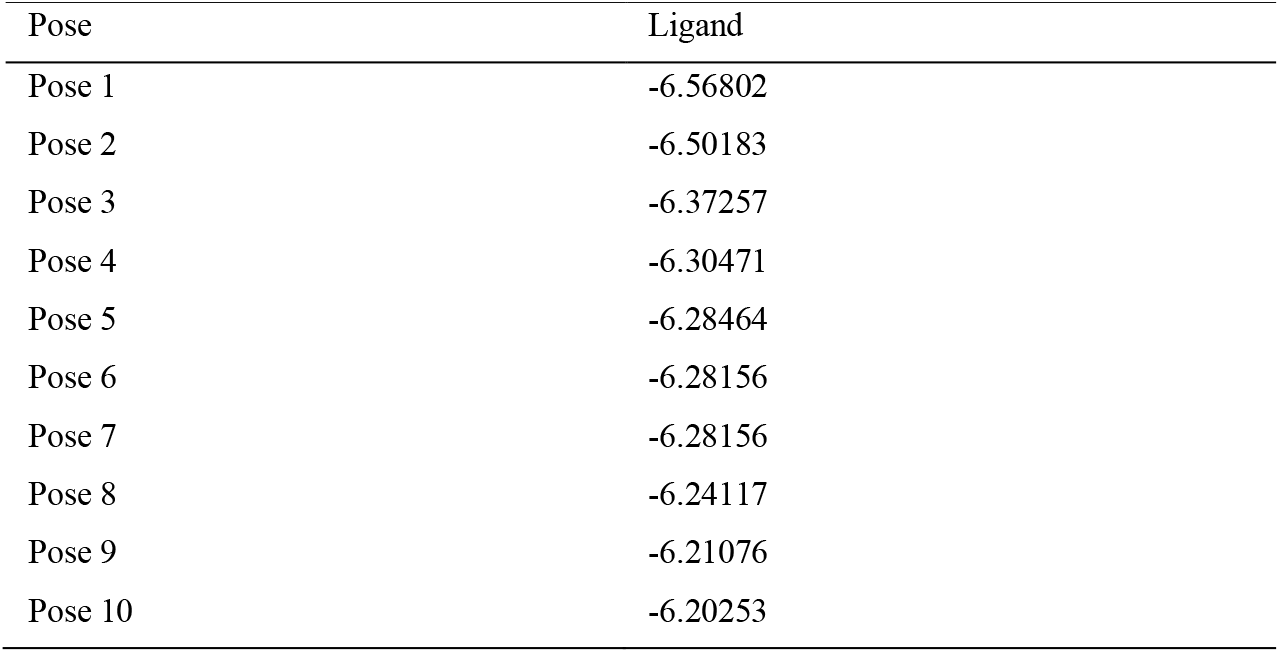
Ligand poses.

**Table 3.**
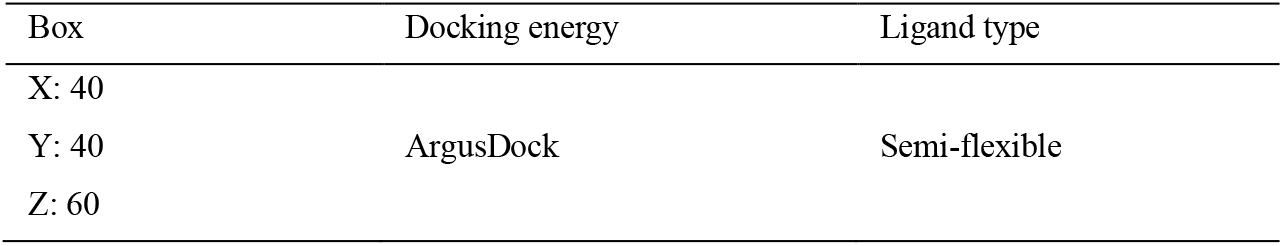
Protein-ligand docking calculation.

| Box | Docking energy | Ligand type |
| --- | --- | --- |
| X: 40 | ArgusDock | Semi-flexible |
| Y: 40 |  |  |
| Z: 60 |  |  |

#### 3.1.2. Conclusions

This part of the study primarily focused on the use of various molecular modeling approaches to predict interactions between vitamin C and staphylococcal FnBPs protein. Molecular docking modeling was performed, followed by the revelation of interactions using Discovery Studio software. The results showed four interactions, both classical and non-classical hydrogen types. These results suggest that vitamin C blocked bacterial entry by binding to its receptor.

### 3.2. Exploration of interactions between vitamin C and the active site of the direct entry receptor gC1qR/p33 of S. aureus

Interactions between vitamin C and the amino acids of the active site were studied, revealing both classical and non-classical hydrogen bonds as well as hydrophobic interactions. Vitamin C exhibited good affinity with the gC1qR/p33 protein, in line with Lipinski’s rule. The study of the interaction between the amino acids of the gC1qR/p33 protein and vitamin C using Arguslab software requires downloading the necessary structures from the databases mentioned earlier, and the structures of the two molecules used in this study are depicted in Figure 4.

**Fig. 4.**
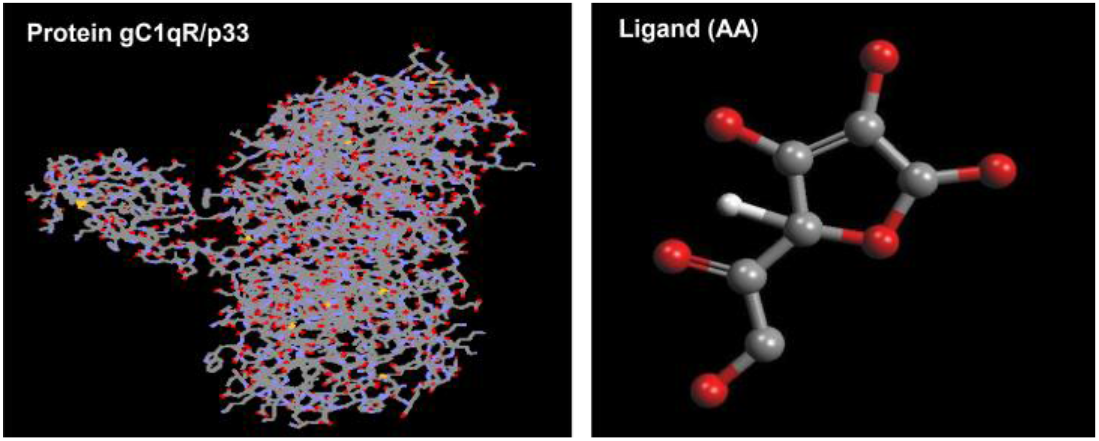
Structure of the gC1qR/p33 protein and the ascorbic acid (AA) ligand.

#### 3.2.1. Exploration of interactions between vitamin C and the active site of the direct entry receptor gC1qR/p33 of S. aureus

The complex formation free energy between ascorbic acid and gC1qR/p33 is obtained using the open-source software ArgusLab (Figure 5).

**Fig. 5.**
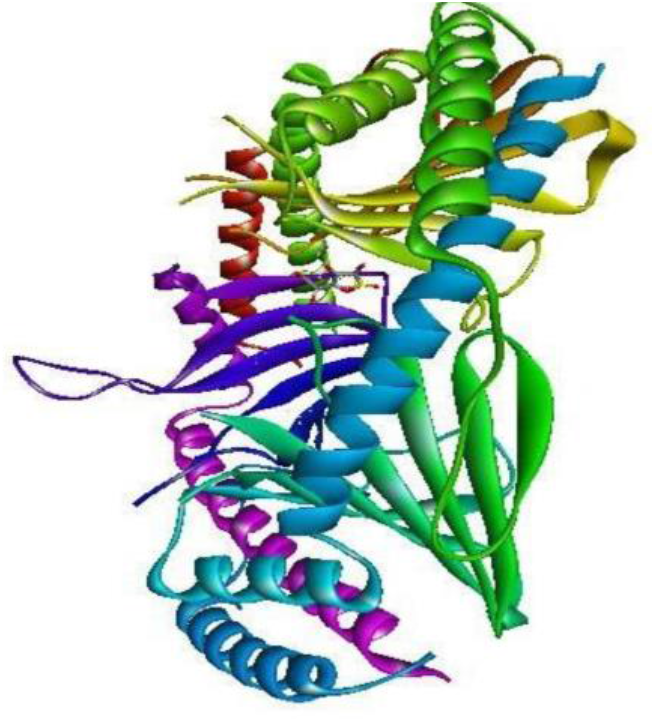
Visualization of the ascorbic acid and gC1qR/p33 molecules.

#### 3.2.2. Molecular docking protein-ligand circulation

The characteristics of ligand-interaction circulation are listed in Table 4.

**Table 4.** Circulation Characteristics (Ligand-Interaction).

| Box | Docking ernaly | Ligand type |
| --- | --- | --- |
| X: 34.000000 | ArgusDock | Semi-flexible |
| Y: 30.000000 |  |  |
| Z: 25.000000 |  |  |

#### 3.2.3. Study of protein-ligand interactions

Molecular docking calculation of interactions between the gC1qR/p33 protein sites and vitamin C revealed an optimal interaction energy with gC1qR/p33 of -5.34639 kcal/mol (*pose 0 fitness), with a docking process runtime of 14 seconds (*Docking run: elapsed time) (Table 5).

**Table 5.** Top 10 molecular docking poses, relative to interactions between the binding sites of the gC1qR/p33 protein and vitamin C.

| Pose | Pose 1 | Pose 2 | Pose 3 | Pose 4 | Pose 5 | Pose 6 | Pose 7 | Pose 8 | Pose 9 | Pose 10 |
| --- | --- | --- | --- | --- | --- | --- | --- | --- | --- | --- |
| Score (Negative Kcal/mol) | 5.34639 | 5.34637 | 5.34621 | 5.19833 | 5.18274 | 5.13107 | 5.12681 | 5.12669 | 5.11563 | 5.11511 |

##### 3.2.3.1. Interactions between ascorbic acid and the active site

Molecular docking allowed the study of interactions between ascorbic acid and the amino acids of the active site of gC1qR/p33 during complex formation. These interactions were ensured through various types of bonds, including classical hydrogen bonds, non-classical hydrogen bonds, and hydrophobic interactions. The list of amino acids involved in the active site and the bonds between ascorbic acid and the amino acids of the active site of gC1qR/p33 during complex formation are illustrated in Figure 6.

**Fig. 6.**
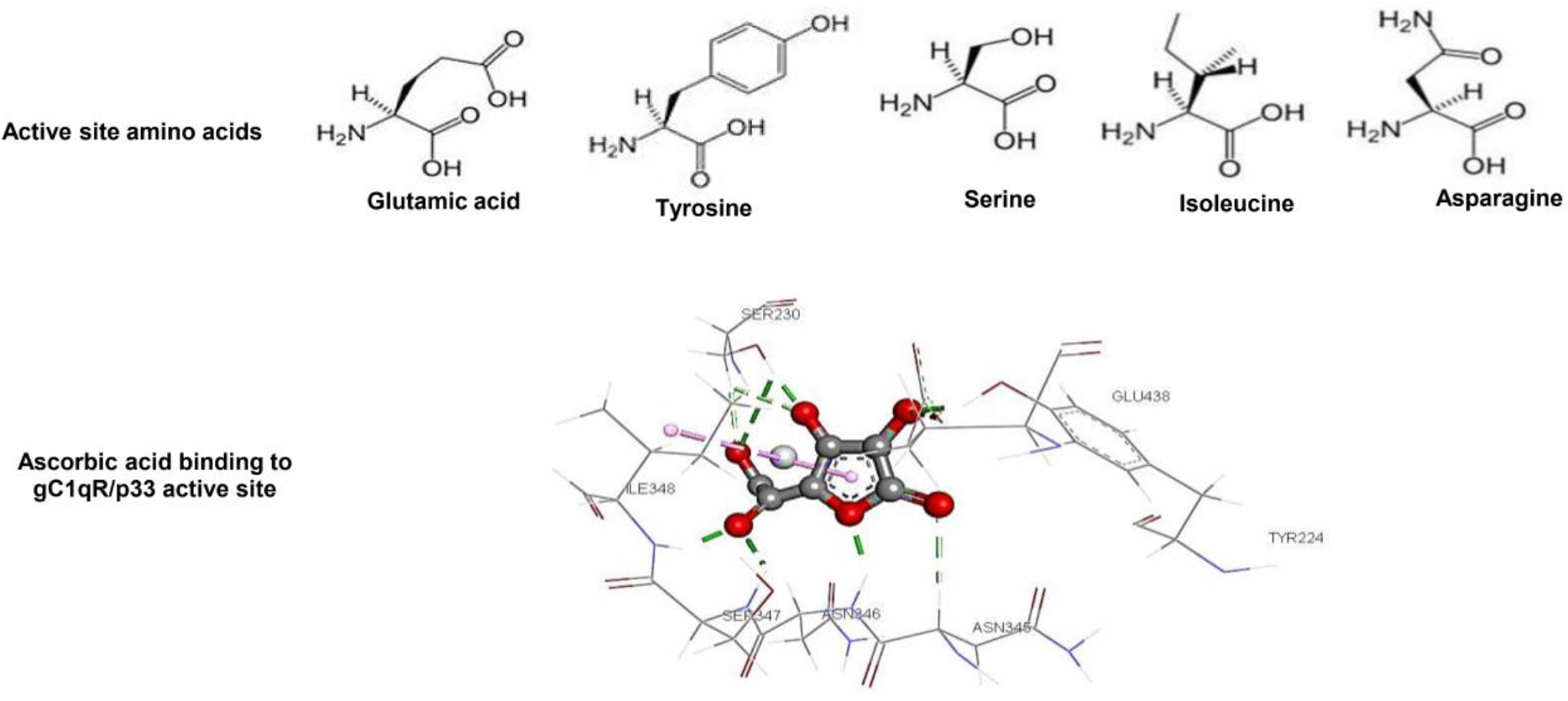
Active site amino acids of gC1qR/p33 and their interactions with ascorbic acid.

##### 3.2.3.2. Hydrogen and hydrophobic interactions

Hydrogen bonding occurs when a hydrogen atom, bound to an electronegative atom (the donor), is attracted to another electronegative atom (the acceptor), acting at very short distances (0.8 to 2.8 Å). These bonds, while few, accommodate well to molecular flexibility. Molecules devoid of charged groups or atoms capable of forming hydrogen bonds are termed hydrophobic substances, as they cannot hydrate and tend to aggregate through coalescence. During docking, ascorbic acid forms six classical hydrogen bonds with six amino acids of the active site of gC1qR/p33: SER230, ILE348, TYR224, ASN346, SER347, and GLU438. Additionally, three non-classical hydrogen bonds are established with two amino acids of the active site: SER230 and ASN345. Molecules lacking charged groups or atoms capable of forming hydrogen bonds are considered hydrophobic and tend to aggregate. A hydrophobic bond is also observed between ascorbic acid and the amino acid ILE348 of the active site, with a distance of 5.028796 A° (Table 6). Classical hydrogen bonds, non-classical hydrogen bonds, and hydrophobic interactions are depicted in Figure 7.

**Table 6.** Hydrogen and Hydrophobic interactions in the protein-ligand complex along with their distances.

| Molecule | Ascorbic acid |  |
| --- | --- | --- |
| Hydrophobic interactions | Involved residues | Distance (Å°) |
|  | ILE348 | 5.028796 |
| Classical hydrogen-bonding interactions | Involved residues | Distance (Å°) |
|  | SER230 | 3.048348 |
|  | GLU438 | 2.848861 |
|  | TYR224 | 1.546703 |
|  | ASN346 | 1.981419 |
|  | SER347 | 2.081011 |
|  | ILE348 | 1.718322 |
| Non-classical hydrogen-bonding interactions | ASN345 | 2.438632 |
|  | SER230 | 2.325086 |
|  | SER230 | 3.052368 |

**Fig. 7.**
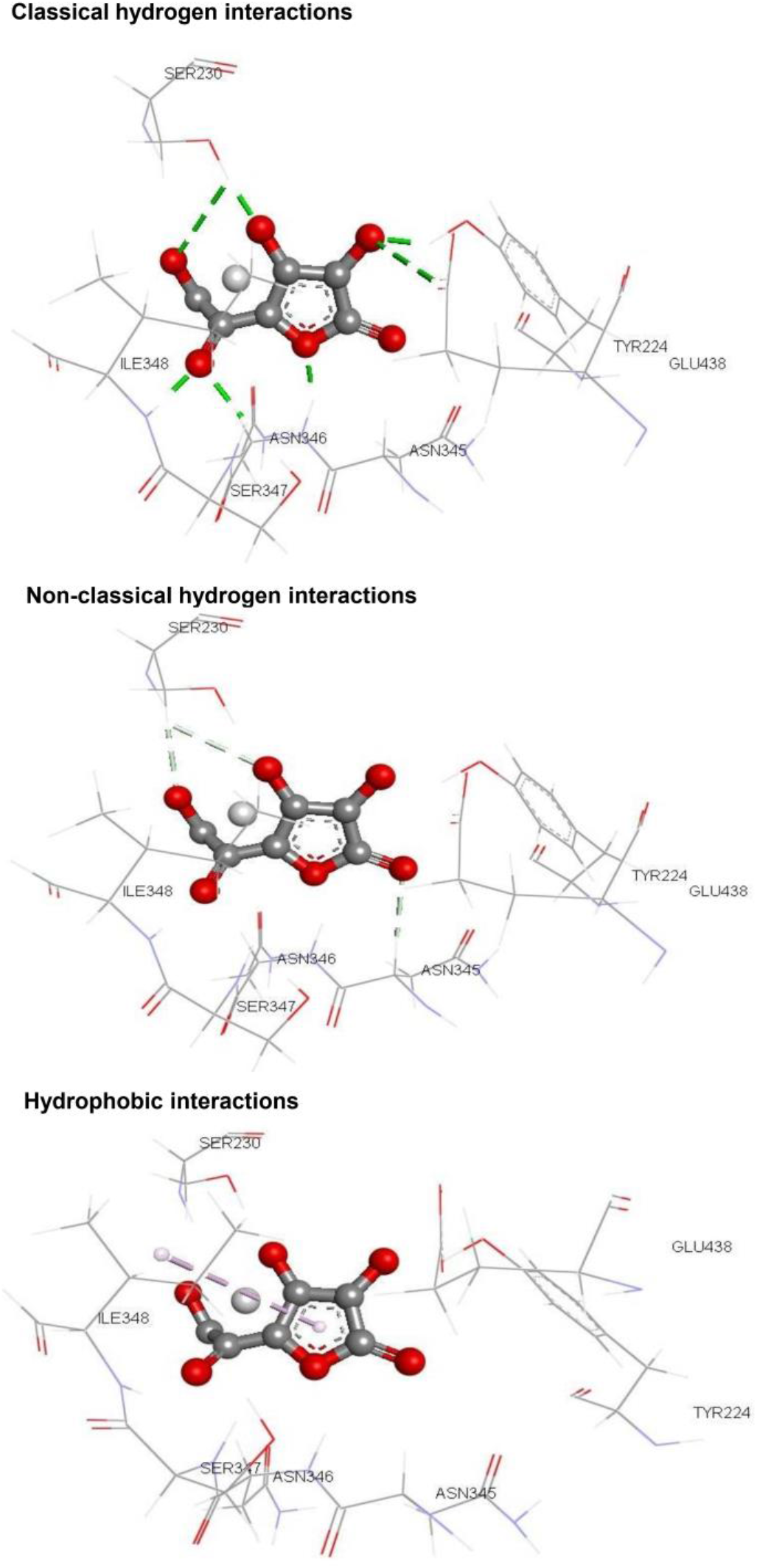
Classic, non-classic hydrogen bonds, and hydrophobic interactions between ascorbic acid and the active site amino acids of gC1qR/p33 during molecular complex formation.

##### 3.2.3.3. Application of Lipinski’s Rule

To assess ligand availability, Lipinski’s rule of 5 was applied. The results of this evaluation are summarized in Table 7. It emerges that vitamin C fully meets Lipinski’s rule criteria. Therefore, vitamin C could be considered a potential inhibitor of the direct entry receptor gc1qr/p33, without presenting absorption issues via oral administration.

**Table 7.** Lipinski rule results on ascorbic acid.

| Compound | MM | logP | HB | HA | FB |
| --- | --- | --- | --- | --- | --- |
| Ascorbic acid | 176.12 | -1.40 | 4 | 6 | 2 |

MM = Molecular Weight ( $\leq 500$ g/mol), LogP = Water/Octanol Partition Coefficient ( $-2 \leq \log P \leq 5$ ), HA = Hydrogen Acceptor (nO, N) ( $\leq 10$ ), HB = Hydrogen Bond Donor (nOH, NH) ( $\leq 5$ ). FB = Flexible Bond (nrotb) ( $\leq 15$ ).

#### 3.2.4. Conclusions

This section focused on the molecular interactions between vitamin C and the gc1qr/p33 direct entry receptor of *S. aureus* in endothelial cells. Analysis of these interactions revealed a specific binding between vitamin C and the active site of gc1qr/p33, involving residues SER230, ASN345, ILE348, TYR224, ASN346, SER347, and GLU438. Additionally, examination of ΔG bind values and application of Lipinski’s rule of 5 confirmed the favorable affinity of vitamin C for the gc1qr/p33 protein. Based on these in silico results, this strong affinity suggests a potential role in preventing endothelial cell infection by *S. aureus*. However, further validation through in vitro and/or in vivo tests may be necessary to confirm these findings.

### 3.3. Study of the effect of vitamin C on staphylococcal protein A

#### 3.3.1. Molecular docking

The active site of the C domain of staphylococcal Protein A and the binding site bounding box are presented in Figure 8. We filled in the dimensions of the binding site bounding box (X = 20, Y = 23, Z = 21) (Table 8). Molecular docking modeling with “ArgusLab” allowed for the reconstruction of the vitamin C ligand, revealing 48 poses with the best score, presenting a ΔG equal to -5.17065 kcal/mol (Table 9). This score results from the formation of hydrogen bonds in the protein’s catalytic cavity (Figure 9). These hydrogen bonds are established with amino acids at distances ranging from 1.661660 to 3.398596 Å. Additionally, numerous hydrophobic interactions were observed, including hydrophobic Pi and Pi/alkyl mixed interactions with distances of 3.557828 Å and 5.322754 Å (Table 10 and Figure 9). These interactions contribute to the stability of the vitamin C - Protein A complex, involving amino acid residues constituting the active site of the protein’s C domain, present on helices 1 (PHE5, GLN9, GLN10, ASN11, PHE13, TYR14, LEU17) and 2 (ASN28, ILE31, GLN32, LYS35).

**Table 8.**
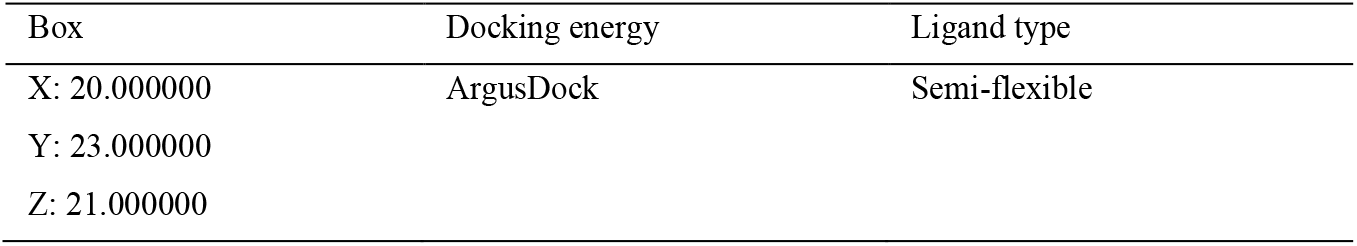
Protein-Ligand Docking Calculation.

**Table 9.**
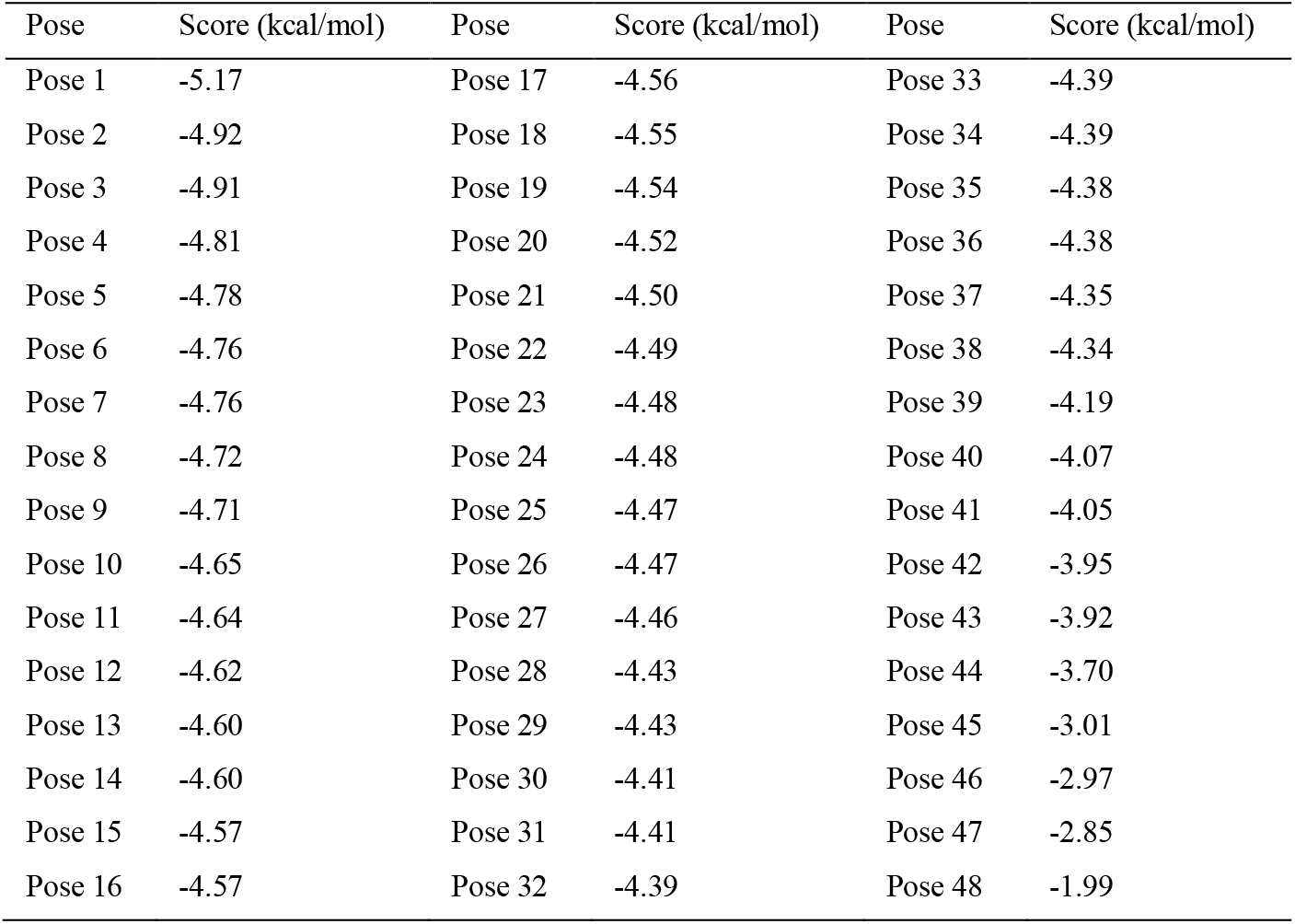
Ligand poses.

| Pose | Score (kcal/mol) | Pose | Score (kcal/mol) | Pose | Score (kcal/mol) |
| --- | --- | --- | --- | --- | --- |
| Pose 1 | -5.17 | Pose 17 | -4.56 | Pose 33 | -4.39 |
| Pose 2 | -4.92 | Pose 18 | -4.55 | Pose 34 | -4.39 |
| Pose 3 | -4.91 | Pose 19 | -4.54 | Pose 35 | -4.38 |
| Pose 4 | -4.81 | Pose 20 | -4.52 | Pose 36 | -4.38 |
| Pose 5 | -4.78 | Pose 21 | -4.50 | Pose 37 | -4.35 |
| Pose 6 | -4.76 | Pose 22 | -4.49 | Pose 38 | -4.34 |
| Pose 7 | -4.76 | Pose 23 | -4.48 | Pose 39 | -4.19 |
| Pose 8 | -4.72 | Pose 24 | -4.48 | Pose 40 | -4.07 |
| Pose 9 | -4.71 | Pose 25 | -4.47 | Pose 41 | -4.05 |
| Pose 10 | -4.65 | Pose 26 | -4.47 | Pose 42 | -3.95 |
| Pose 11 | -4.64 | Pose 27 | -4.46 | Pose 43 | -3.92 |
| Pose 12 | -4.62 | Pose 28 | -4.43 | Pose 44 | -3.70 |
| Pose 13 | -4.60 | Pose 29 | -4.43 | Pose 45 | -3.01 |
| Pose 14 | -4.60 | Pose 30 | -4.41 | Pose 46 | -2.97 |
| Pose 15 | -4.57 | Pose 31 | -4.41 | Pose 47 | -2.85 |
| Pose 16 | -4.57 | Pose 32 | -4.39 | Pose 48 | -1.99 |

**Table 10.** Types of Bonds Established between the Protein and the Ligand.

| Molecule | Hydrogen interaction |  | hydrophobic interaction |  |
| --- | --- | --- | --- | --- |
|  | Involved residues | Distance (Å) | Involved residues | Distance (Å) |
| Ascorbic acid | PHE13 | 3.398596 | PHE13 | 5.322754 |
|  | GLN9: HE22 | 2.208768 | LYS35 | 3.557828 |
|  | GLN9: HE21 | 2.897263 |  |  |
|  | ASP2 | 1.661660 |  |  |

**Fig. 8.**
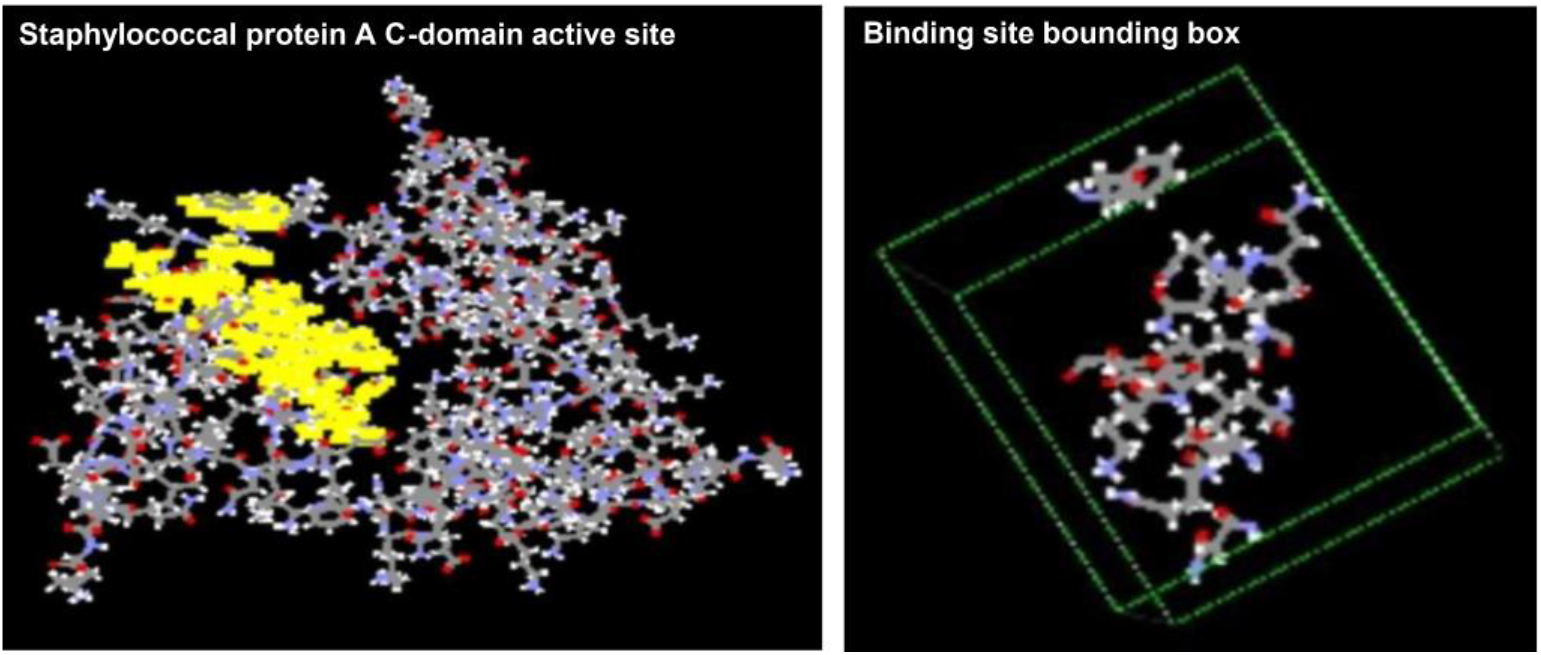
Active site of the C domain of the staphylococcal Protein A and the site bounding box. We specified the dimensions of the site bounding box (X = 20, Y = 23, Z = 21).

**Fig. 9.**
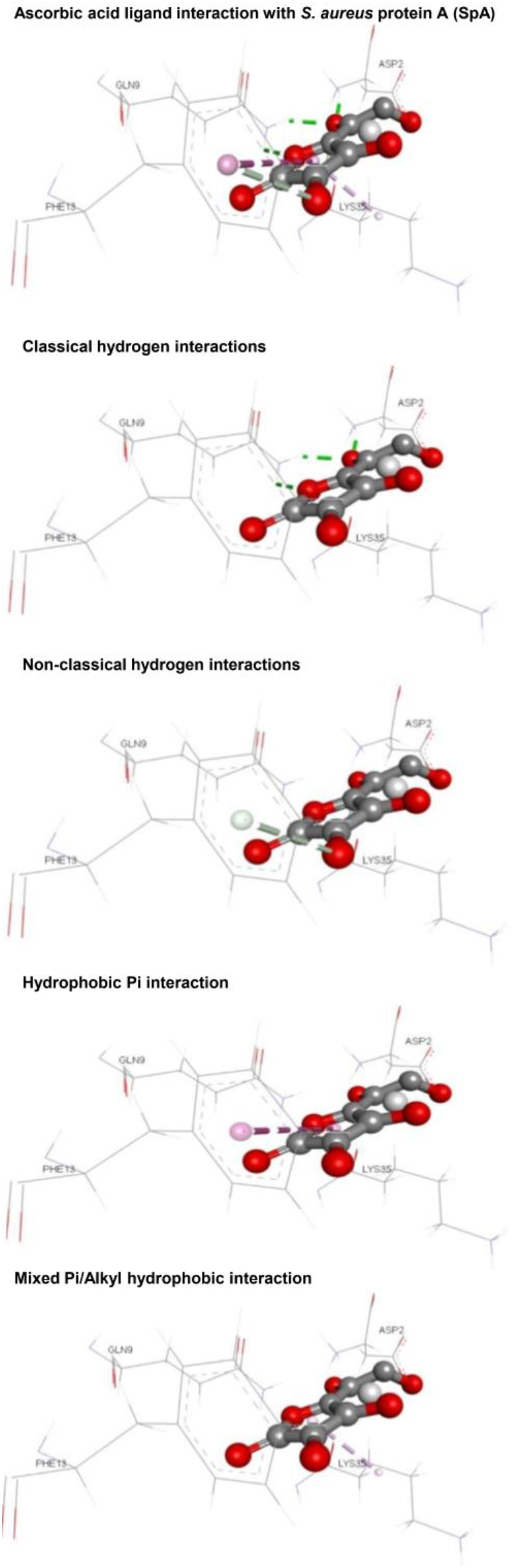
Interactions of the ligand with the protein, including hydrogen bonds, non-classic hydrogen bonds, hydrophobic Pi interactions, and mixed Pi/alkyl hydrophobic interactions.

#### 3.3.2. Interaction between vitamin C and Protein A

Complex 4WWI, displaying the lowest energy score (−5.17065 kcal/mol) (Table 8), established 6 interactions with our ligand. Hydrogen bonds are formed with the following amino acids: PHE13 at a distance of 3.4 Å, GLN9: HE22 at a distance of 2.2 Å, GLN9: HE21 at a distance of 2.8 Å, as well as with ASP2 at a distance of 1.6 Å. It is also noteworthy to mention the presence of hydrophobic interactions with amino acid LYS35 at a distance of 3.5 Å, as well as a second interaction with PHE13 at a distance of 5.3 Å (Table 9). These interactions suggest that vitamin C could have an impact on staphylococcal Protein A.

#### 3.3.3. Lipinski’s rule of five

Each ligand must meet several fundamental criteria, including affordable production cost, adequate solubility and stability, as well as absorption, distribution, metabolism, excretion, and toxicity characteristics. In this regard, we evaluated the bioavailability of our ligand using the Molinspiration Checker. This evaluation was based on the criteria established by Lipinski’s rules. The evaluated parameters include a molecular weight below 500 daltons, a partition coefficient (logP) or lipophilicity between -2 and 5, a maximum of five hydrogen bond donors, a maximum of ten hydrogen bond acceptors, and a number of rotation angles less than or equal to five. Our results show that our ligand fully satisfies Lipinski’s criteria, which bodes well for good interaction even without ligand optimization. The results of applying these rules to our ligand are presented in Table 11

**Table 11.** Lipinski’s Rules application on vitamin C.

| Ligand | LogP | PSA | Natcoms | MW | nON | Nohnh | nviolation | nrot |
| --- | --- | --- | --- | --- | --- | --- | --- | --- |
| Ascorbic acid | -1.40 | 107.22 | 12 | 176.12 | 6 | 4 | 0 | 2 |

#### 3.3.4. Conclusions

This study primarily focused on the use of several theoretical molecular modeling approaches to predict interactions between vitamin C and staphylococcal Protein A. Molecular docking modeling was performed, followed by the revelation of interactions using Discovery Studio software. The six identified interactions included both classical and non-classical hydrogen bonds, as well as Pi and mixed Pi/alkyl hydrophobic interactions. These results suggest that vitamin C could have an inhibitory effect on Protein A, a virulence factor of the bacterium *S. aureus*.

## 4. Conclusions and Future Prospects

It is evident to conclude that vitamin C has strong potential to interfere with interactions between *S. aureus* and endothelial cells. This conclusion arises from the results obtained from various molecular modeling approaches, including molecular docking, which revealed specific molecular interactions between vitamin C and *S. aureus* cell receptors. Thus, vitamin C specifically interacts with staphylococcal FnBPs and α5β1 proteins, as well as with the active sites of target proteins, including the gC1qR/p33 direct entry receptor and staphylococcal Protein A, thereby inhibiting *S. aureus* adhesion and entry into endothelial cells (Table 12). These interactions involve different residues, confirming a favorable affinity of vitamin C for these proteins. Indeed, in the study on gC1qR/p33, classical and non-classical hydrogen bonds were observed with key residues such as SER230, ASN345, ILE348, TYR224, ASN346, SER347, and GLU438, as well as hydrophobic interactions, demonstrating diversified interaction with the active site. Similarly, in the study on staphylococcal Protein A, hydrophobic interactions of Pi and mixed Pi/alkyl types were identified, highlighting the complexity of the interaction between vitamin C and this protein. This affinity could play a crucial role in preventing endothelial infections by *S. aureus*. This affinity was corroborated by the analysis of ΔG bind values and the conformity of vitamin C to Lipinski’s rule of 5 (Table 13), which evaluates the bioavailability of pharmaceutical compounds. Molecular modeling studies have identified active sites and key residues involved in these interactions, thus highlighting the potential mechanism by which vitamin C exerts its preventive effect against bacterial infections.

**Table 12.**
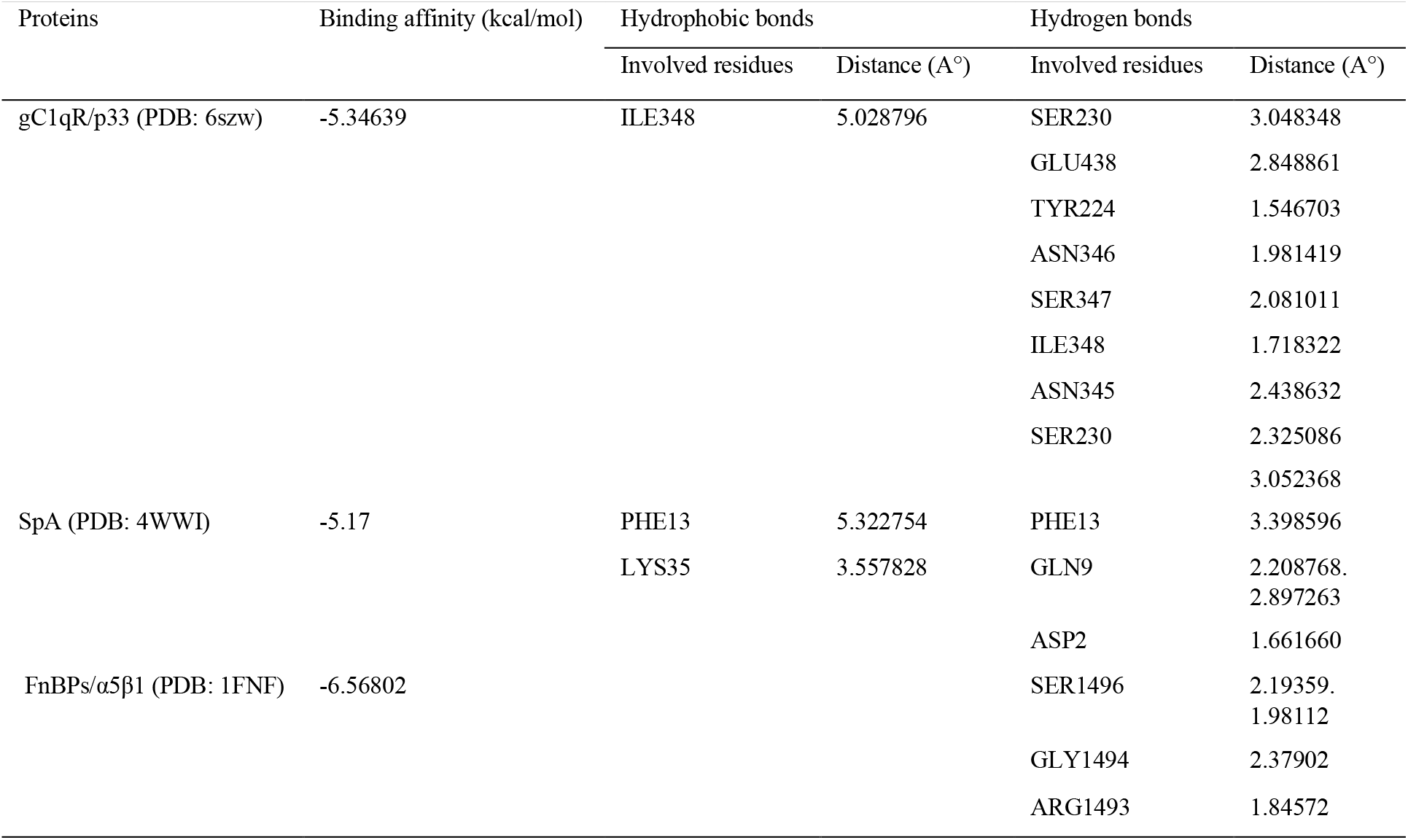
Molecular docking score. interactions and bond length of gC1qR/p33, SpA and FnBPs/α5β1 with vitamin C.

**Table 13.** Physicochemical properties and Lipinski’s Rule of five.

|  | LogP | HBD | HBA | MW | LRO5M met |
| --- | --- | --- | --- | --- | --- |
| Rule | < 5 | ≤ 5 | ≤ 10 | < 500 | At least 3 |
| Ascorbic acid | -1.40 | 4 | 6 | 176.12 | 2 |

HBA: hydrogen bond acceptors, HBD: hydrogen bond donors, LogP: logarithm of the partition coefficient (P) between n-octanol and water, LRO5M: Lipinski's Rule of five, MW: molecular weight.

In terms of prospects, it is important to note that these results require further validation through in vitro and/or in vivo experiments to fully confirm their clinical relevance. Further studies could also explore the underlying molecular mechanisms of vitamin C action and its potential efficacy as a therapeutic agent against bacterial infections. These researches could include studies on bacterial resistance mechanisms to vitamin C and the identification of synergies with other antimicrobial agents. These results could pave the way for new strategies for the prevention and treatment of infectious diseases, thus offering new perspectives in the fields of public health and medicine.

## Conflict of Interest Statement

The authors have no conflicts of interest to declare.

## Acknowledgments

The authors extend their gratitude to the Ministry of Higher Education and Scientific Research for their support of this work under the PRFU projects, with registration ID D01N01UN130120200004.

## Notes

### Competing Interest Statement

The authors have declared no competing interest.

### Summary of Updates

Staphylococcus aureus (S. aureus), often perceived as an extracellular pathogenic agent, exhibits a remarkable ability to penetrate host cells, including endothelial, epithelial, and osteoblastic cells, significantly contributing to the pathogenesis of infections. A significant pathway for this invasion appears to involve the bacterium's binding to the α5β1 integrin via a fibronectin bridge, followed by phagocytosis. Additionally, S. aureus presents staphylococcal protein A, a cell wall protein that binds to the Fc and Fab regions of immunoglobulins, playing a crucial role in virulence and immune evasion, and can also bind directly to the gC1qR receptor on endothelial cells. Furthermore, vitamin C is recognized for its antimicrobial and immunomodulatory properties, offering potential for reducing the risk of infection. Considering the aforementioned elements, our study focused on exploring the potential effects of vitamin C on the interactions between S. aureus and endothelial cells. Thus, we particularly examined two aspects: on the one hand, interactions involving fibronectin-binding proteins (FnBPs) proteins and human α5β1 integrin, and on the other hand, the interaction between vitamin C and the direct entry receptor (the globular heads of complement component 1q receptor [gC1qR/p33], also known as hyaluronic acid binding protein 1 [HABP1]). To achieve this, we utilized molecular modeling assays, primarily relying on molecular docking.

## References

[1] E.I.B. Peerschke, B. Ghebrehiwet, The contribution of gC1qR/p33 in infection and inflammation, Immunobiology 212 (2007) 333–342. 10.1016/j.imbio.2006.11.011.

[2] A.C. Carr, S. Maggini, Vitamin C and Immune Function, Nutrients 9 (2017) 1211. 10.3390/nu9111211.

[3] S. Chambial, S. Dwivedi, K.K. Shukla, P.J. John, P. Sharma, Vitamin C in disease prevention and cure: an overview, Indian J. Clin. Biochem. IJCB 28 (2013) 314–328. 10.1007/s12291-013-0375-3.

[4] S. Mousavi, S. Bereswill, M.M. Heimesaat, Immunomodulatory and Antimicrobial Effects of Vitamin C, Eur. J. Microbiol. Immunol. 9 (2019) 73–79. 10.1556/1886.2019.00016.

[5] J. Kallio, M. Jaakkola, M. Mäki, P. Kilpeläinen, V. Virtanen, Vitamin C inhibits staphylococcus aureus growth and enhances the inhibitory effect of quercetin on growth of Escherichia coli in vitro, Planta Med. 78 (2012) 1824–1830. 10.1055/s-0032-1315388.

[6] S. Dahou, M.C.-E. Smahi, W. Nouari, Z. Dahmani, S. Benmansour, L. Ysmail-Dahlouk, M. Miliani, F. Yebdri, N. Fakir, M.Y. Laoufi, M. Chaib-Draa, A. Tourabi, M. Aribi, L-Threoascorbic acid treatment promotes S. aureus-infected primary human endothelial cells survival and function, as well as intracellular bacterial killing, and immunomodulates the release of IL-1β and soluble ICAM-1, Int. Immunopharmacol. 95 (2021) 107476. 10.1016/j.intimp.2021.107476.

[7] J.A. Tuszynski, P. Winter, D. White, C.-Y. Tseng, K.K. Sahu, F. Gentile, I. Spasevska, S.I. Omar, N. Nayebi, C.D. Churchill, M. Klobukowski, R.M.A. El-Magd, Mathematical and computational modeling in biology at multiple scales, Theor. Biol. Med. Model. 11 (2014) 52. 10.1186/1742-4682-11-52.

[8] D.A. Antunes, C.T. Schoeder, M. Baek, E.A. Donadi, Editorial: Structural modeling and computational analyses of immune system molecules, Front. Immunol. 14 (2023) 1274670. 10.3389/fimmu.2023.1274670.

[9] S.A. Hollingsworth, R.O. Dror, Molecular Dynamics Simulation for All, Neuron 99 (2018) 1129–1143. 10.1016/j.neuron.2018.08.011.

[10] L.H.S. Santos, R.S. Ferreira, E.R. Caffarena, Integrating Molecular Docking and Molecular Dynamics Simulations, Methods Mol. Biol. Clifton NJ 2053 (2019) 13–34. 10.1007/978-1-4939-9752-7_2.

[11] S.K. Burley, H.M. Berman, G.J. Kleywegt, J.L. Markley, H. Nakamura, S. Velankar, Protein Data Bank (PDB): The Single Global Macromolecular Structure Archive, Methods Mol. Biol. Clifton NJ 1607 (2017) 627–641. 10.1007/978-1-4939-7000-1_26.

[12] S. Kim, T. Cheng, S. He, P.A. Thiessen, Q. Li, A. Gindulyte, E.E. Bolton, PubChem Protein, Gene, Pathway, and Taxonomy Data Collections: Bridging Biology and Chemistry through Target-Centric Views of PubChem Data, J. Mol. Biol. 434 (2022) 167514. 10.1016/j.jmb.2022.167514.

[13] G.R. Sridhar, P.V. Nageswara Rao, D.S. Kaladhar, T.U. Devi, S.V. Kumar, In Silico Docking of HNF-1a Receptor Ligands, Adv. Bioinforma. 2012 (2012) 705435. 10.1155/2012/705435.

[14] D. Iqbal, M. Alsaweed, Q.M.S. Jamal, M.R. Asad, S.M.D. Rizvi, M.R. Rizvi, H.M. Albadrani, M. Hamed, S. Jahan, H. Alyenbaawi, Pharmacophore-Based Screening, Molecular Docking, and Dynamic Simulation of Fungal Metabolites as Inhibitors of Multi-Targets in Neurodegenerative Disorders, Biomolecules 13 (2023) 1613. 10.3390/biom13111613.

